# Heterogeneity across mammalian- and avian-origin A(H1N1) influenza viruses influences viral infectivity following incubation with host bacteria from the human respiratory tract

**DOI:** 10.1101/2025.03.28.645935

**Authors:** Poulami Basu Thakur, Hannah A. Bullock, James Stevens, Amrita Kumar, Taronna R. Maines, Jessica A. Belser

**Author notes:** Correspondence: Influenza Division, 1600 Clifton Rd NE, MS H17-5, Atlanta, GA, 30329.

## Abstract

Influenza A viruses (IAV) are primarily transmitted between mammals by the respiratory route, and encounter bacteria in the respiratory tract before infecting susceptible epithelial cells. Previous studies have shown that mammalian-origin IAV can bind to the surface of different bacterial species and purified bacterial lipopolysaccharides (LPS), but despite the broad host range of IAV, few studies have included avian-origin IAV in these assessments. Since IAV that circulate in humans and birds are well-adapted to replication in the human respiratory and avian gastrointestinal tracts, respectively, we investigated the ability of multiple human and avian A(H1N1) IAV to associate with bacteria and their surface components isolated from both host niches. Binding interactions were assessed with microbial glycan microarrays, revealing that seasonal and avian IAV strains exhibited binding diversity to multiple bacterial glycans at the level of the virus and the bacterium, independent of sialic acid binding preference of the virus. Co-incubation of diverse IAV with LPS derived from *Pseudomonas aeruginosa* (*P. aeruginosa*), a respiratory tract bacterium, led to reduced retention of viral infectivity in a temperature dependent manner which was not observed when co-incubated with LPS from *Escherichia coli*, a gut bacterial isolate. Reduction of viral infectivity was supported by disruption of IAV virions following incubation with *P. aeruginosa* LPS using electron microscopy. Our findings highlight that both human and avian IAV can bind to bacterial surface components from different host sites resulting in differential functional interactions early after binding, suggesting the need to study IAV-bacteria interactions at the host range interface.

## Introduction

Influenza A viruses (IAV) have a broad host range and infect a diversity of animal species in addition to causing annual seasonal infections in human populations. Disease presentation of seasonal influenza virus infections in humans can vary from asymptomatic to fatal depending on both viral- and host-specific factors. Estimates of the annual disease burden of seasonal influenza are around one billion cases, with three to five million cases of severe disease, and 290,000 to 650,000 respiratory deaths worldwide (1). IAV can also cause rare pandemics due to zoonotic IAV spillover events that result in successful infection of human hosts and sustained human-to-human transmission. Such outbreaks may result in increased patient morbidity and mortality due to a lack of pre-existing immunity within human populations to most zoonotic viruses (2). However, documented cases of human infection with zoonotic IAV are typically self-limiting without onward spread to susceptible contacts, due to a variety of viral and host determinants that restrict virus host range (3, 4).

Seasonal and pandemic influenza virus infections are often followed by serious and potentially fatal respiratory complications caused by secondary bacterial infections (5–7). These bacterial infections can present during acute viral infection, synergizing with the virus to exacerbate disease incidence, or after clearance of the virus from the host by the immune system (8). Although the respiratory tract is home to a wide diversity of bacteria, the most frequently occurring pathogens associated with secondary bacterial infections include *Streptococcus pneumoniae*, *Staphylococcus aureus* and *Haemophilus influenzae* (8, 9). Multiple studies have shown that the pathogenesis of different bacterial species in the human respiratory tract can be enhanced by prior IAV infection (5, 6, 10, 11). These studies involving interactions between IAV and host bacteria have largely employed human-adapted seasonal IAV, which preferentially bind α-2,6 linked sialic acid (SA) receptors prevalent on the surface of epithelial cells in the human respiratory tract. Zoonotic IAV that arise from non-human hosts, however, can differ from their human counterparts in their preferred sites of replication as well as their routes of transmission. Most avian IAV, for example, preferentially bind α-2,3 linked SA receptors present on epithelial cells of the avian gut and are primarily transmitted by the fecal-oral route between avian species. Furthermore, avian viruses are well adapted to replicate at higher temperatures of the avian gut compared to the cooler temperatures of the human upper respiratory tract (URT). Prior studies have evaluated the relative contribution of many viral and host properties towards efficient replication of IAV at distinct host sites of replication (4). However, these studies have largely overlooked host niche-specific bacterial populations, encountered by human and avian IAV, prior to reaching susceptible epithelial cells across different host species and at different tissues within the same host species.

Most previous research assessing IAV-bacteria interactions has investigated the consequence of prior IAV infection on subsequent bacterial pathogenicity, with a paucity of studies assessing these interactions’ capacity to modulate IAV infectivity. Bandoro and Runstadler (12) observed a reduction in stability of an A(H1N1) strain of IAV after incubation with a lipopolysaccharide (LPS) derived from *E. coli*, with data supporting that this decrease in virion stability was associated with morphological changes to the virion structure of a lab-adapted A(H1N1) strain due to incubation with the LPS. Another study by Rowe et al. (13) showed that removal of bacteria from the ferret respiratory tract, using an antibiotic applied topically to the nostrils, abolished airborne transmission of a seasonal IAV in this species, with restoration of virus transmissibility following experimental colonization of the ferret nostril with *S. pneumoniae*. However, while some studies have included side-by-side comparisons of human and avian IAV (12), there remains a critical knowledge gap in understanding the nature of interactions between bacteria and zoonotic IAV, and if interactions between zoonotic IAV and bacterial populations in the mammalian URT represent a host range determinant (4). Our study aims to address that gap by assessing how seasonal and avian IAV interact with host bacteria, present at different host sites, that support virus replication and how these interactions impact virus infectivity. Understanding interactions between host bacteria and both seasonal and zoonotic IAVs will improve our assessment of the relative threat that these viruses pose to human health.

## Methods

### Viruses in study

A(H1N1)pdm09 viruses derived from the 2009 pandemic, A/Nebraska/14/2019 and A/Nebraska/15/2018, were propagated in Madin-Darby canine kidney (MDCK) cells placed in a humidified cell culture incubator with 95% O_2_ and 5% CO_2_. Avian A(H1N1) viruses, A/duck/New York/15024-21/1996 and A/ruddy turnstone/Delaware/300/2009, were propagated in MDCK cells or 10- to 11-day-old embryonated hens’ eggs, respectively, as previously described (14). Stock titers of all viruses were obtained by standard plaque assay in MDCK cells as previously described (15, 16). All experiments with avian viruses were conducted under biosafety level 2 containment, including enhancements (BSL-2E) as required by the U.S. Department of Agriculture.

### Amino acid residues in viral HA

Viral HA protein sequences were obtained from GISAID or NCBI. Amino acid sequences were aligned using BioEdit Sequence Alignment Editor version 7.2.

### Replication kinetics of IAV in Calu-3 cells

The human bronchial epithelial cell line Calu-3 (ATCC) was cultured as previously described (16). Cells were grown to confluence under submerged conditions in 12 well plates with transwell inserts (Corning) at 37°C in a humidified cell culture incubator with 95% O_2_ and 5% CO_2._ Prior to inoculation, apical media was removed from the cell monolayer and cells were washed to remove serum present in culture medium. Cells were infected apically in triplicate with 300 μL of IAV at a multiplicity of infection (MOI) of 0.01 for one hour and then washed and incubated at 33°C or 37°C with 500 µL apical media to emulate temperatures in the URT and lower respiratory tract (LRT), respectively. Infected cell supernatants were collected apically at 2, 24, 48 and 72 hours post infection and immediately frozen at −80°C until titration. Virus titers were assessed in cell supernatants by standard plaque assay in MDCK cells. Growth curves were generated and analyzed using Prism 6.0.7 (GraphPad Software Inc).

### Assessment of interaction between IAV and bacteria

We assessed the binding/interaction between IAV and wildtype *P. aeruginosa* serotype 10 (ATCC). Briefly, *P. aeruginosa* was sub-cultured (1:50) in 25 mL of lysogeny broth (LB) from a 5 mL overnight culture in LB and allowed to grow to mid-log phase (OD_600_ ∼ 0.5-0.6) with shaking at 37°C. The culture was then washed in PBS and adjusted to a concentration of 10^8^ colony forming units (CFU) per ml in PBS. IAV were diluted to 10^6^ plaque forming units (PFU) per ml in PBS. 1 mL of IAV was mixed with 1 mL of *P. aeruginosa* and incubated at 33°C for 2 hours. Co-incubated samples were centrifuged for 10 mins at 5,000 x *g*. The pellet and supernatant fractions were collected and treated with 1X Penicillin/Streptomycin (Gibco) and immediately frozen by transferring to -80°C. Samples were stored at -80°C until further processing. Infectivity readouts were obtained for all IAV using standard plaque assay technique with MDCK cells. Statistical significance between IAV incubated in PBS and IAV incubated with *P. aeruginosa* was obtained by Student’s t-test or by one-way ANOVA for each virus from experiments four independent replicates.

### Inactivation, purification and labeling of IAV

Stock viruses diluted in serum free medium were used to infect MDCK cells with TPCK (tosyl phenylalanyl chloromethyl ketone) trypsin. Viruses were harvested 48 hours after infection and HA titers were obtained for each virus. IAV were inactivated by adding 50 µL of β-propiolactone dropwise to 50 mL of cell supernatant containing the viruses and passaged twice in eggs to ensure their innocuity. Following confirmation of innocuity, IAV were purified through 25% sucrose in NTE buffer (100 mM NaCl, 10 mM Tris, 1 mM EDTA) using ultracentrifugation for 2 hours at 28,000 rpm. IAV were resuspended in PBS overnight and centrifuged again at 15,000 rpm for 1 hour to remove any residual sucrose. IAV were then resuspended in PBS and HA titers were obtained. Viral HA titers were adjusted to 256 hemagglutination units (HAU) per 50 µL. Purified IAV were then fluorescently labeled by mixing with 1 M NaHCO_3_ (pH 9.0) and Alexa Fluor 488 dye (Thermo Fisher Scientific) and incubating labeled viruses at room temperature for 1 hour in the dark. Labeled IAV were then dialyzed in one liter of PBS for 1 hour by gentle rotation of dialysis membrane units (Thermo Fisher Scientific) at 4°C. The dialysis buffer was replaced with fresh PBS after an hour and IAV were allowed to dialyze further overnight at 4°C. The dialysis buffer was replaced again following overnight dialysis and IAV were dialyzed for 1 hour at 4°C. IAV were then transferred from dialysis units to 2 mL screw-cap tubes and stored at 4°C until further processing.

### Assessment of binding of IAV to sialic acids and bacterial polysaccharides

We assessed the binding of IAV to different sialic acids and to a diversity of purified bacterial polysaccharides in collaboration with the Emory University Glycomics Core. The Legacy Consortium for Functional Glycomics (CFG) glycan array, version 6, was used to assess the binding preference of IAV to different sialic acid-containing glycans. The Microbial Glycan Microarray (MGM), version 2 was used to assess binding interactions between IAV and polysaccharides purified from different bacterial species. For both arrays, 70 µL of labeled IAV at 256 HAU per 50 µL in PBS was spiked with 0.7 µL 10% (w/v) BSA and 0.7 µL 5% (v/v) Tween 20. Samples were applied to the Legacy CFG v6, array to assess viruses binding to sialic acid containing glycans, and to the MGM array to evaluate binding to bacterial polysaccharides. Slides were incubated at room temperature in a dark humidified chamber for 1 hour then washed four times with PBS containing 0.05% (v/v) Tween 20, four times with PBS and then four times with water. Slides were then subjected to spin drying and then scanned with InnoScan 1100AL scanner (resolution: 5 µm/pixel, 488 nm laser: PMT 85/Laser Power High). Data processing was performed using the Mapix 8.6.0 software (Innopsys, Chicago, IL). Data were processed by the Emory University Glycomics Core to obtain basic descriptive statistics. No normalization method was used.

### Visualization of binding of IAV to bacterial LPS

Stock IAV were diluted two-fold. 100 µL of diluted IAV (minimum concentration of 1.3e5 PFU/100µl) were co-incubated with 100 µg LPS derived from *P. aeruginosa* serotype 10 or *E. coli* O111:B4 (Sigma Aldrich), or with 100 µL of water for 1 hour at 33°C. Diluted IAV incubated for one hour at 4°C was used as a temperature control. Control or co-incubated IAV were visualized using transmission electron microscopy (TEM) in collaboration with the Infectious Diseases Pathology Branch at CDC. Specimens for TEM were fixed in 2.5% paraformaldehyde, incubated on nickel TEM grids for two hours, and stained with 5% ammonium molybdate (pH 6.9) with 0.1% trehalose (17). EM grids were visualized using a FEI/Thermo Fisher Tecnai Spirit electron microscope.

### Evaluation of cellular internalization of IAV following incubation with bacterial LPS

MDCK cells were cultured on glass coverslips (Thermo Scientific) in 24 well plates (Corning) at 37°C for 24 hours. Cells were washed with PBS twice and replenished with serum free growth medium. Our panel of IAV was diluted to an MOI of 2; 100 µL of diluted IAV was incubated with 100 µL PBS or with 100 µL of PBS containing 100 µg LPS derived from *P. aeruginosa* and *E.coli*, for 2 hours at 33°C. 100 µL of IAV alone or co-incubated with LPS was used to infect MDCK cells at 37°C for one hour. Following infection, the inoculum was removed, and the cells were washed with PBS and replenished with serum-free growth medium. At 8 hours post infection, the cells were fixed with 3.7% paraformaldehyde and then permeabilized with 0.25% (v/v) Triton X-100 (Sigma Aldrich) for 30 minutes. Permeabilized cells were incubated with mouse anti-nucleoprotein (NP) monoclonal antibody A-3 (25), followed by Alexa Fluor 488-conjugated secondary antibody (Thermo Fisher Scientific). Immunostained cells were counterstained with 4′,6-diamidino-2-phenyindole (DAPI) (Sigma-Aldrich) for nuclei and examined under a Zeiss LSM710 confocal fluorescence microscope (Carl Zeiss Microscopy, LLC). After counting of DAPI- and NP-positive cells in 4 independent 40× fields of view, the infection rate was determined as the number of NP-positive cells per 100 cells in each field.

### Coincubation of IAV with bacterial polysaccharides to assess retention of viral infectivity

Stock IAV was diluted to 10^6^ PFU/mL in PBS unless otherwise specified. 100 µL of diluted IAV was incubated with 100 µL of 1 mg/mL of bacterial lipopolysaccharides (LPS) derived from *P. aeruginosa* or *E. coli* and with water or PBS for two hours at 33°C or 42°C to emulate temperatures of the mammalian URT and the avian gut, respectively. Following incubation, 100 µL of co-incubated virus was titrated on MDCK cells. IAV diluted in PBS was incubated for the duration of the assay at 4°C and used as a control. Infectivity was measured using IAV titers obtained from standard plaque assay. The percentage of retention of infectivity remaining after incubation with PBS or LPS at different temperatures was calculated by dividing the titer of each sample by that of the control sample. Results are shown from three independent experiments.

## Results

### Seasonal and avian H1N1 IAV exhibit diversity in binding to sialic acid receptors

Seasonal and avian IAV replicate in distinct host sites by binding to different sialylated glycans on the surface of susceptible and permissive host cells, with human adapted seasonal IAV preferentially binding to α-2,6 linked sialic acids and avian IAV preferentially binding to α-2,3 linked sialic acids (18, 19). As IAV have been shown to bind sialic acids present on the bacterial surface (20, 21), we first determined the receptor binding specificity of a panel of seasonal and avian A(H1N1) viruses. Two contemporary circulating A(H1N1)pdm09 influenza A viruses, A/Nebraska/14/2019 (Neb/14) and A/Nebraska/15/2018 (Neb/15), and two avian A(H1N1) viruses, A/duck/New York/15024-21/1996 (dk/NY) and A/ruddy turnstone/Delaware/300/2009 (rd/Ts), were chosen in this study to encompass human-adapted (Neb/14 and Neb/15), land-based poultry (dk/NY), and wild shorebird (rd/Ts) strains. Neb/14 and Neb/15 viruses share high sequence identity with original A(H1N1)pdm09 isolates including the well-studied A/California/07/2009 and were chosen to represent viruses that had exhibited sustained human spread for many years. The human strains, Neb/14 and Neb/15, differ at residue 190 in the viral hemagglutinin (HA) (Table 1) that plays a role in receptor binding (22) and are predicted to bind α-2-6 linked sialic acids. The avian isolates have the avian consensus 190E/225G predictive of α-2-3 linked sialic acid binding (23, 24).

**Table 1.**
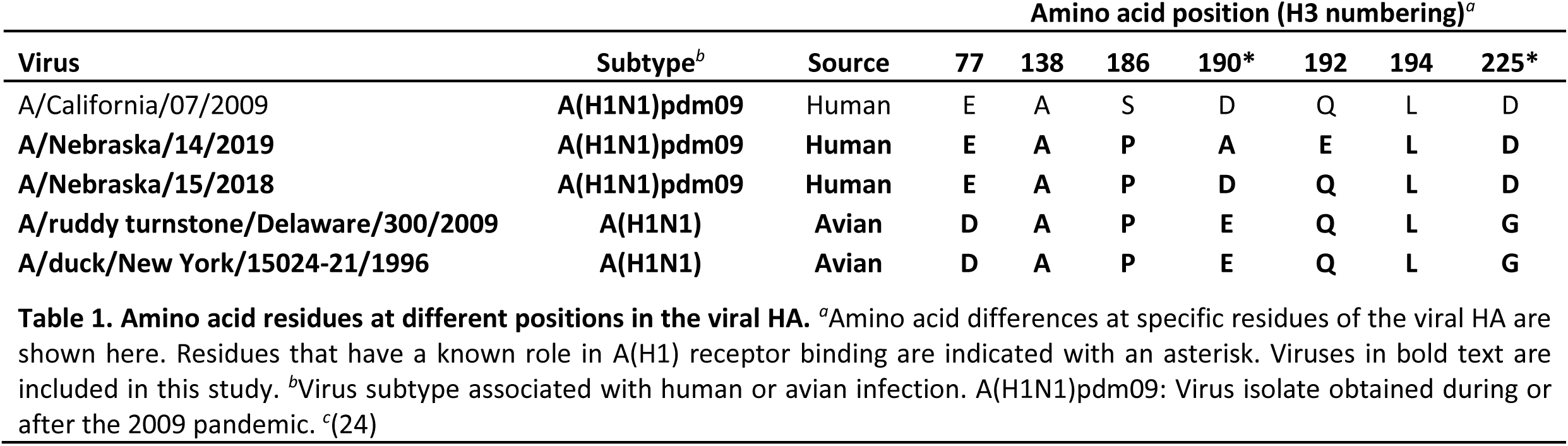
Amino acid residues at different positions in the viral HA. *^a^*Amino acid differences at specific residues of the viral HA are shown here. Residues that have a known role in A(H1) receptor binding are indicated with an asterisk. Viruses in bold text are included in this study. *^b^*Virus subtype associated with human or avian infection. A(H1N1)pdm09: Virus isolate obtained during or after the 2009 pandemic. *^c^*(24)

| Virus | Subtype<sup>b</sup> | Source | Amino acid position (H3 numbering)<sup>a</sup> |  |  |  |  |  |  |
| --- | --- | --- | --- | --- | --- | --- | --- | --- | --- |
|  |  |  | 77 | 138 | 186 | 190* | 192 | 194 | 225* |
| A/California/07/2009 | **A(H1N1)pdm09** | Human | E | A | S | D | Q | L | D |
| **A/Nebraska/14/2019** | **A(H1N1)pdm09** | **Human** | **E** | **A** | **P** | **A** | **E** | **L** | **D** |
| **A/Nebraska/15/2018** | **A(H1N1)pdm09** | **Human** | **E** | **A** | **P** | **D** | **Q** | **L** | **D** |
| **A/ruddy turnstone/Delaware/300/2009** | **A(H1N1)** | **Avian** | **D** | **A** | **P** | **E** | **Q** | **L** | **G** |
| **A/duck/New York/15024-21/1996** | **A(H1N1)** | **Avian** | **D** | **A** | **P** | **E** | **Q** | **L** | **G** |

To determine receptor binding specificity, both seasonal and avian IAV were subjected to glycan microarray screening. As expected, seasonal IAV showed a strong binding preference to α-2,6 linked sialic acids and minimal binding to α-2,3 linked sialic acids on the array (Figure 1a, b). The avian strain dk/NY showed a strong binding preference to α-2,3 linked sialic acids and weak binding preference for α-2,6 linked sialic acids (Figure 1c), previously shown by Van Hoeven et al (25). In contrast, the avian strain rd/Ts exhibited a mixed binding preference to both α-2,3 and α-2,6 linked sialic acids (Figure 1d), in agreement with previous work by Koçer et al (26).

**Figure 1.**
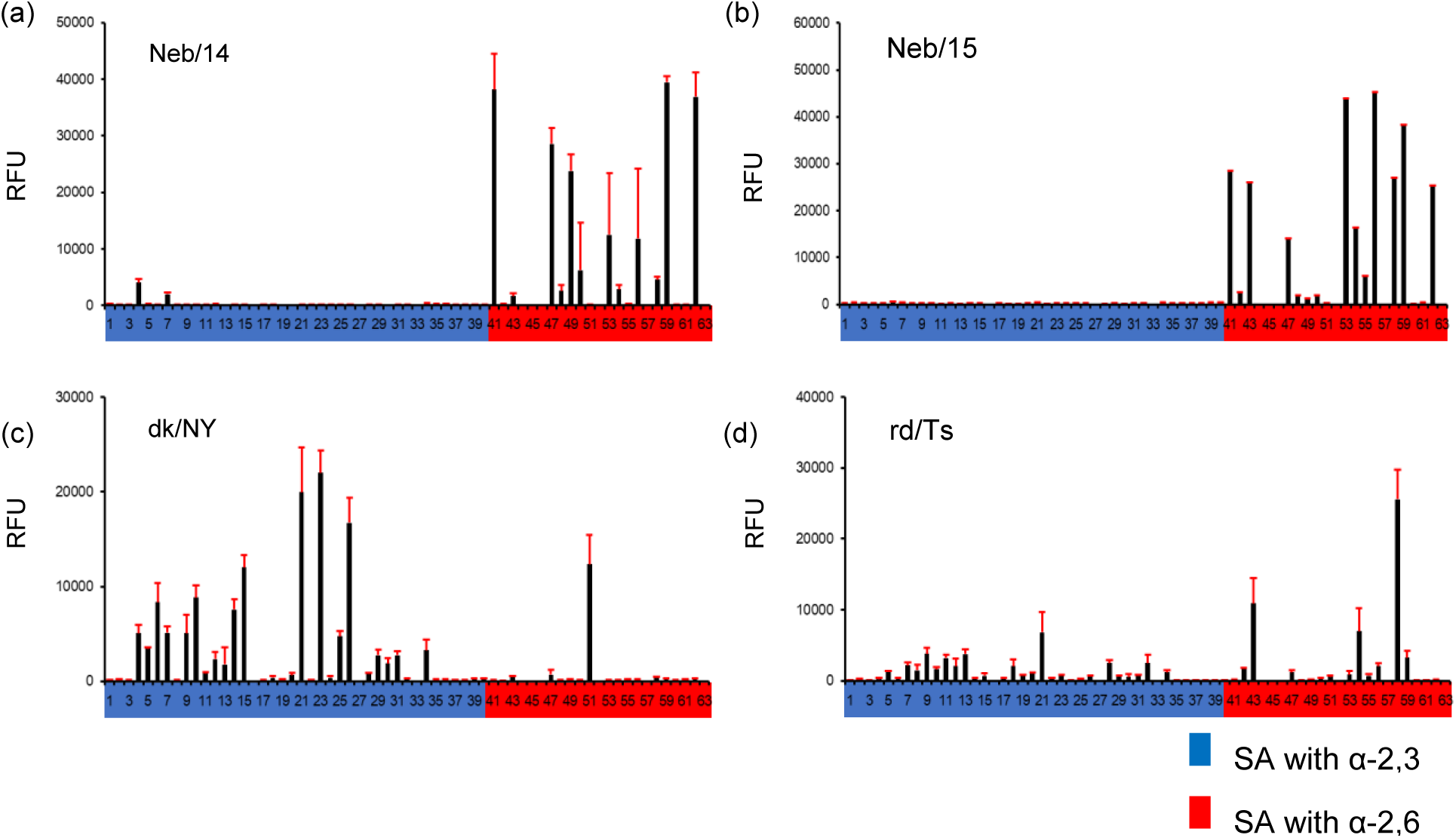
Glycan microarray-based binding specificities of human and avian H1N1 IAV. Glycan binding preference of contemporary circulating human H1N1 IAV, Neb/14 **(a)** and Neb/15 **(b)**, avian A(H1N1) IAV, dk/NY **(c)** and rd/Ts **(d)** are presented as evaluated by fluorescence intensity of binding to the array. Colored bars indicate α-2,3 linked sialic acids (blue) and α-2,6 linked sialic acids (red). Error bars represent the standard deviation (SD) for 4 independent replicates on the array. RFU, relative fluorescence units.

### Seasonal and avian influenza viruses are capable of replicating in human bronchial epithelial cells

To assess the replicative capacity of avian A(H1N1) IAV in the human airway relative to seasonal IAV, we evaluated the ability of our panel of influenza A viruses to replicate in Calu-3 human bronchial epithelial cells at 33°C and 37°C, temperatures of the URT and LRT, respectively. All viruses tested were able to replicate productively in Calu-3 cells at both temperatures when inoculated at a multiplicity of infection (MOI) of 0.01 (Figure 2). As expected, both seasonal viruses replicated efficiently at both temperatures, reaching mean peak titers at 48 hours post infection (hpi). Surprisingly, the LPAI virus dk/NY was able to reach mean titers comparable to the seasonal IAV at all timepoints examined, while rd/Ts yielded significantly lower titers at 48 and 72 hpi at both temperatures (p<0.005) compared to the other viruses. However, despite observing lower infectious titers of the rd/Ts strain, both avian A(H1N1) viruses tested were able to replicate at 33°C and 37°C, indicating the capacity of these strains to replicate at temperatures found throughout the human respiratory tract.

**Figure 2.**
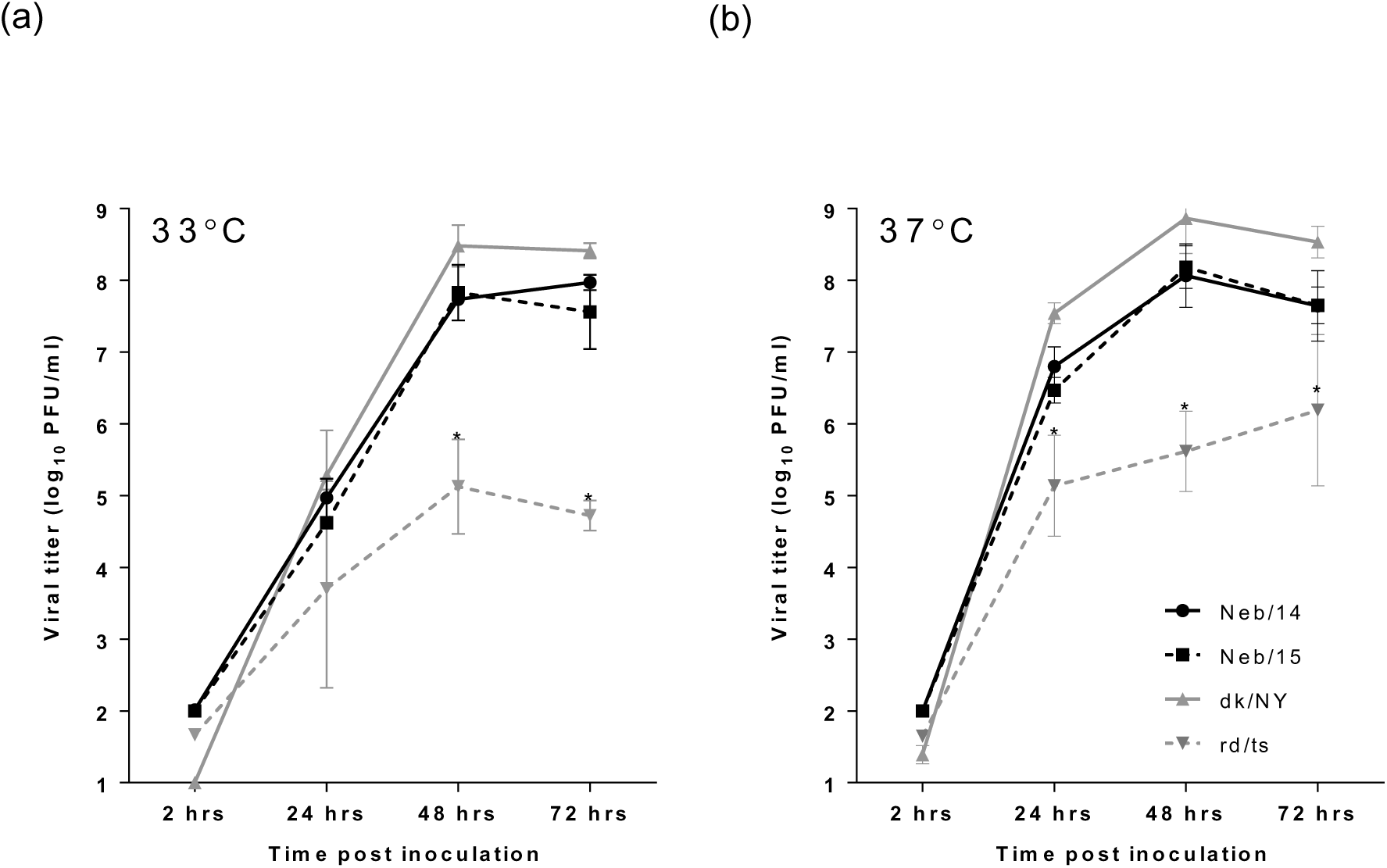
Replication of IAV in Calu-3 cells. Calu-3 cells grown in liquid culture were infected with human and avian IAV at a MOI 0.01 and cultured at **(a)** 33°C or **(b)** 37°C. Culture supernatants were collected at the indicated times post inoculation. Virus titers were determined by standard plaque assay in MDCK cells to quantify infectious virus. The limit of detection was 10 PFU. Bars indicate mean of three independent replicates and standard deviation. Statistical significance between viral titers at each time point was determined by two-way analysis of variance (ANOVA). * p < 0.005.

### IAVs co-sediment with *P. aeruginosa* after incubation

Our viral replication kinetics data suggest that avian A(H1N1) can maintain strain-specific replication fitness at the cooler airway temperatures of the mammalian URT. As the URT hosts different bacterial species than the LRT or the avian GI tract (27), we next examined if human and avian IAV could associate with bacteria from the mammalian URT and if such associations adversely impacted viral infectivity. To emulate conditions encountered by the virus in the human URT, both human and avian IAV were incubated at 33°C with either *P. aeruginosa* or PBS as a control. We compared infectious titers of IAVs incubated in PBS to those incubated with *P. aeruginosa* to assess if incubation with the bacterium would compromise infectivity of the virus. No significant loss of infectivity of either human or avian IAV strains was observed when incubated with *P. aeruginosa* when compared to incubation in PBS only (Figure 3a). To quantify the measure of association between IAVs and *P. aeruginosa* we used a modified co-sedimentation assay (10) where IAVs incubated with the bacterium were centrifuged and infectious virus titers were assessed in the pellet and supernatant fractions. Our results indicate that both human and avian IAV strains were able to co-sediment with the bacterium, though the percentage of infectious virus recovered in the pellet fraction varied by virus. The human viruses, Neb/14 and Neb/15, showed 35.1 and 59.7 percent of total infectious titers in the pellet post-incubation, and 64.9 and 40.3 percent infectious titers in the supernatant, respectively. Contrastingly, the avian strains retained 67.8 (dk/NY) and 16.0 (rd/Ts) percent infectious titers in the pellet, and 32.2 and 84.0 percent infectious titers in the supernatant, respectively. Thus, our data suggest that avian IAVs could associate with a bacterial species of the human URT as with the human strains. However, the nature of such associations was strain specific for both human and avian viruses.

**Figure 3.**
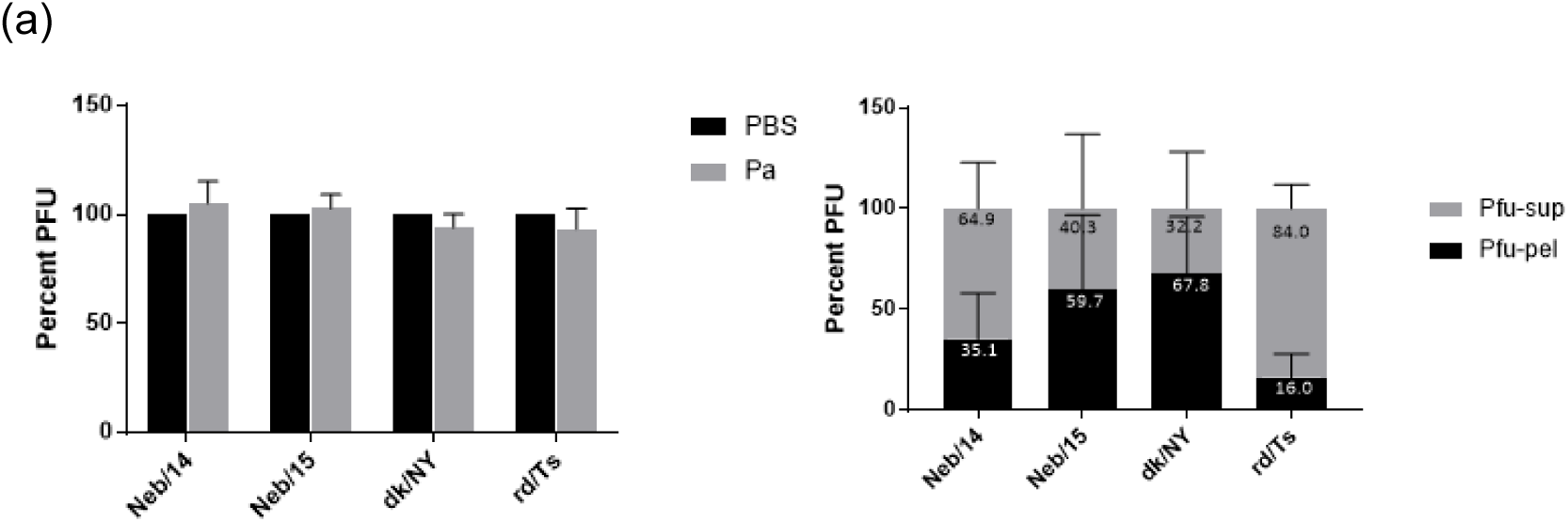
Co-sedimentation of IAV with *P. aeruginosa*. Human and avian IAV were incubated at 33°C in PBS or with *P. aeruginosa* for 2 hours prior to centrifugation, with titration of pellet and supernatant fractions. (a) Infectious titers from pellet and supernatant fractions combined were assessed for control and *P. aeruginosa* treated IAVs using a standard plaque assay in MDCK cells. (b) The percentage of infectious virus recovered from the *P. aeruginosa* pellet and supernatant fractions following titration by standard plaque assay with MDCK cells. Error bars indicate mean of four independent replicates and standard deviation.

### IAVs bind to diverse bacterial glycans independent of sialic acid binding preference

To further explore the complexity of interkingdom interactions between A(H1N1) IAV and microbial populations, and to identify with more accuracy the breadth of specific bacterial targets bound by IAV, we conducted subsequent studies using bacterial polysaccharides. The microbial glycan microarray (28) contains 313 purified polysaccharides from both prokaryotic and eukaryotic species. Of these, 308 targets are derived from a diversity of bacterial species including, but not limited to, those present in the human respiratory tract and extrapulmonary sites such as the GI tract. This array has been employed previously to assess interactions between host immune factors and diverse microbes, though its use in assessing binding interactions against whole virus has not been previously included in virology studies. To assess if IAV with distinct SA binding preferences exhibited differential binding to bacterial polysaccharides, human and avian IAV were inactivated and labeled with a fluorophore and subjected to the MGM array. As viruses were standardized by HA titers across glycan microarrays (and not intensity of the fluorescent signal), direct comparisons of relative binding strength to specific binding targets between different viruses were not possible. However, by ranking binding order strength (and not relative fluorescence units (RFU)) within microbial targets bound to a specific IAV, we could identify patterns in binding present across different IAV.

All IAV tested exhibited binding to bacterial polysaccharides on the microarray, though strain-specific differences in binding patterns were detected between the human and avian IAV examined. The seasonal A(H1N1)pdm09 IAVs Neb/14 and Neb/15 (which showed a strong binding preference to α-2,6 linked sialic acids (Figure 1a,b)), exhibited greater commonality of binding to bacterial polysaccharides within the top 20% of polysaccharides in the array (n=63) as compared to dk/NY and rd/Ts avian IAVs (Figure 4a-i, ii), among which more variability was detected. Interestingly, top binding hits of the α-2,3 binding dk/NY virus substantially overlapped with glycans bound by both human viruses (Figure 4a-iii), in contrast to the dual-binding rd/Ts virus which showed lower commonality of binding to these targets (Figure 4a-iv). Top hits for both human IAV, Neb/14 and Neb/15 as well as the avian virus, dk/NY, included surface polysaccharides from *Yersinia pestis* and *Salmonella typhimurium.* Interestingly, rd/Ts differed from the other viruses by binding to glycans from *Proteus* spp. and *Shigella flexneri* (Figure 4c). Similar results were observed in the binding of human and avian IAVs to microbial glycans within the top 10% and 30% of glycans on the array (data not shown), collectively supporting that IAV from different hosts are able to bind to a diversity of bacterial glycans independent of their sialic acid binding preference.

**Figure 4.**
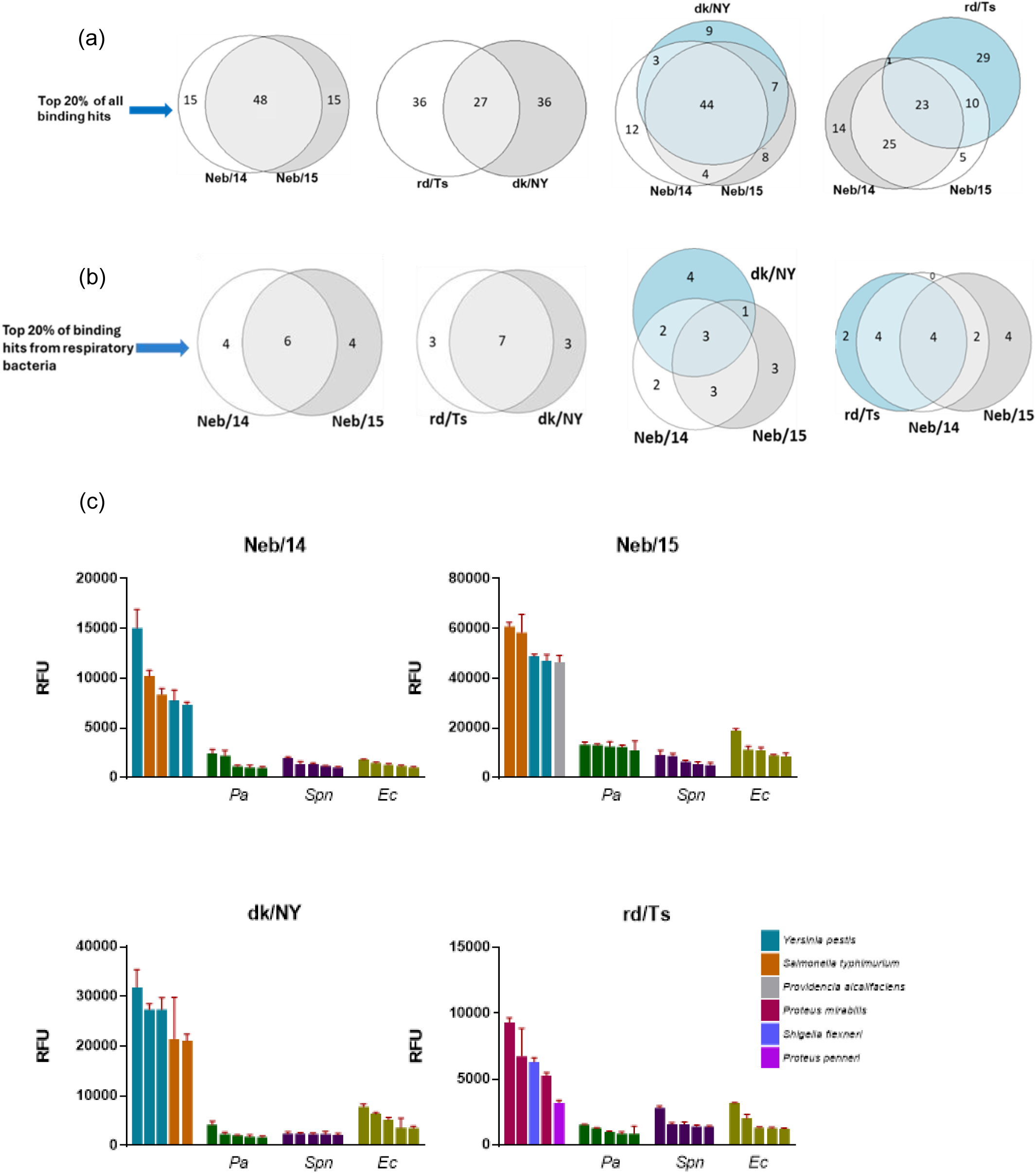
Microbial glycan microarray based binding interactions between IAV and bacterial polysaccharides. Microbial glycan binding commonality of human and avian IAV within the (a) top 20% (n=63) of all bacterial polysaccharide targets and the (b) top 20% (n=10) of polysaccharides from respiratory bacterial species are indicated here. (c) The highest intensity of binding of IAV to the top five targets on the microarray are indicated on the left side of each graph, and relative binding of IAV to the top five glycans derived from each bacterium specified that inhabit the respiratory or GI tract are shown here. *Pa*: *P. aeruginosa*, *Spn*: *S. pneumoniae*, *Ec*: *E. coli*

Next, we wanted to assess whether different host-origin IAV strains showed differential binding to glycans derived specifically from bacterial species found in the respiratory tract. For all IAV assessed, relative binding to glycans derived from bacteria that inhabit the respiratory tract was substantially lower in comparison to binding to the top glycan targets on the microarray (Figure 4c). When restricted to the top 20% of glycans of respiratory bacteria on the array (n=10), both human and avian viruses showed a high degree of binding to the same targets, with generally comparable overlap between all four IAV strains tested (Figure 4b). These results suggest a high degree of conservation among binding to targets IAV may encounter in the human airways despite different levels of mammalian host adaptation of the viruses themselves.

Assessments of relative binding strength of IAV to different glycans from the same bacterial species are not frequently performed. Thus, microarrays can serve as high throughput tools that permit concurrent assessment of relative strength of binding between IAV and glycans from different bacterial strains within the same species. We ranked all included targets for *P. aeruginosa* and *E. coli*, representing species frequently detected in the human respiratory tract and GI tract, to the human virus Neb/14, and compared the order of relative binding strength to all the other viruses tested. Even within the same bacterial species, diversity of binding affinity to glycans was observed for each IAV (Figure 5). Strain specific variation was noted in binding to the same glycans between seasonal and avian IAV, and within the same host origin IAV. Taken together, all IAV bound most strongly on each array to non-respiratory bacteria derived products, suggesting that IAV-bacterial binding events detected in the array were driven by the structural features presented on bacterial glycans, and not other factors that may be associated with the specific bacterial species with which they interacted. Among respiratory tract bacterial derived products, binding was observed to a range of these glycans for all IAV. These data support that whole IAV can bind to a diversity of bacterial products, including those present at sites of IAV replication in the mammalian respiratory tract, with differences in binding strength varying within products derived from the same bacterial species.

**Figure 5.**
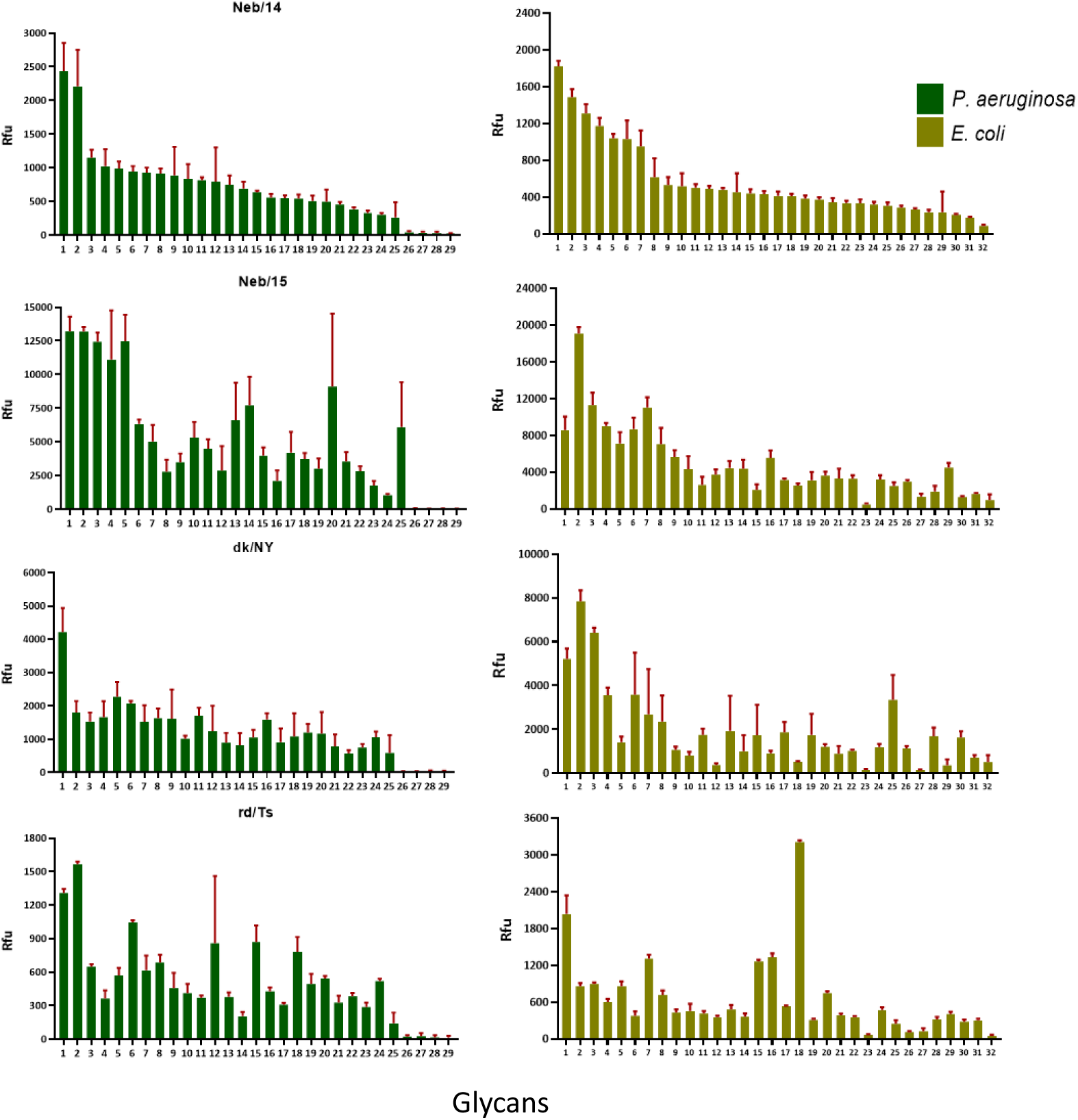
Relative binding strength of IAV to different glycans from bacterial species of the respiratory and GI tract. Binding of the human virus, Neb/14 to glycans derived from isolates of *P. aeruginosa* and *E. coli* were ranked according to the intensity of the signal and compared to the relative binding strength of all other IAV. Glycans from different species of *P. aeruginosa* and *E. coli* were ordered from the highest to the lowest binding signal on the array for the human IAV strain Neb/14. The order of bacterial glycans that all other IAV strains bound to were standardized to the order of glycans that Neb/14 bound to. Glycan 5 derived from a *P. aeruginosa* serotype 10 and glycan 7 derived from *E. coli*, strain O111: B4 were used in subsequent assays.

### Incubation with lipopolysaccharide from *P. aeruginosa* causes disruption of the virion envelope

Following detection of binding between IAV and diverse bacterial surface glycans, we next assessed if there was a functional consequence to the virus after binding to LPS. Since IAV is exposed to the bacterial milieu in the mammalian URT before infecting cells in this specific host niche, we incubated Neb/14, a human virus, with LPS isolated from *P. aeruginosa*, a respiratory bacterial pathogen, at 33°C. Results were visualized using transmission electron microscopy (TEM) with a magnification of 26,000x (Figure 6). We observed binding between Neb/14 to *P. aeruginosa* derived LPS within 1 hour of incubation. Greater disruption of the virion envelope was observed with incubation of IAV with LPS from *P. aeruginosa* in comparison to the virus incubated at 4°C as a control, with elevated disruption detected rapidly (within 5 minutes) post-incubation, suggesting LPS derived from *P. aeruginosa* binds to and disrupts the envelope of IAV virions as visualized by TEM.

**Figure 6.**
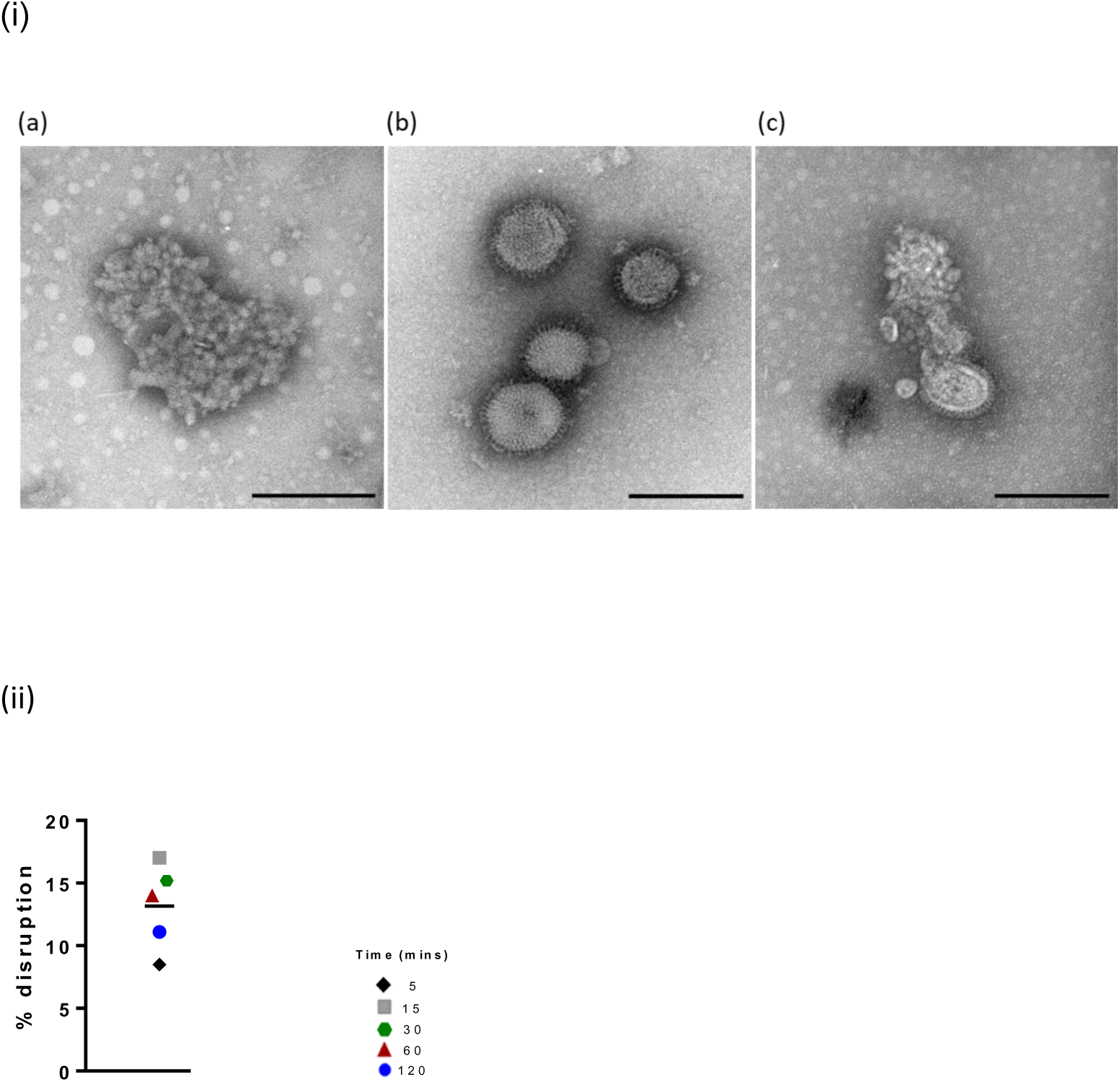
Binding to, and disruption of the viral envelope following coincubation with LPS. (i) TEM micrographs of isolated LPS from *P. aeruginosa* (a), Neb/14 in PBS after incubation for 1 hour at 4°C (b), Neb/14 with LPS from *P. aeruginosa* after incubation for 1 hour at 33°C (c) are shown. IAV were imaged using negative stain electron microscopy. Representative TEM micrographs were obtained at a magnification of 26,000x. Scale bars are 200 nm. (ii) LPS derived from *P. aeruginosa* was incubated with IAV for varying lengths of time. Percent disruption of Neb/14 virions without exposure to LPS and following incubation with *P. aeruginosa* LPS at different times are indicated. n/a: stock virus diluted in PBS only with no disruption observed.

### Lipopolysaccharide derived from *P. aeruginosa* reduces infectivity of seasonal and avian influenza viruses

Since we observed increased rapid disruption of virion envelopes upon incubation of IAV with LPS from *P. aeruginosa*, we next wanted to assess if this disruption in IAV envelopes could impact the ability of these viruses to initiate infection. We incubated the human virus Neb/14 with LPS derived from *P. aeruginosa* and *E. coli* at 33°C, temperatures emulative of the human URT. MDCK cells were then infected with IAV co-incubated with LPS following which the inocula were removed and cells were fixed 7 hours later. Fluorescently stained viral NP merged onto DAPI stained cellular nuclei were qualitatively assessed for infection (Fig. 7). We observed lower levels of infection in cells infected with Neb/14 virus incubated with LPS from *P. aeruginosa* compared to cells infected with the virus incubated in PBS or with LPS from *E. coli* as assessed by the presence of NP+ cells. Our results suggest that LPS derived from *P. aeruginosa* not only causes disruption of the virion envelope as assessed by TEM, but also decreased levels of infection in MDCK cells.

**Figure 7.**
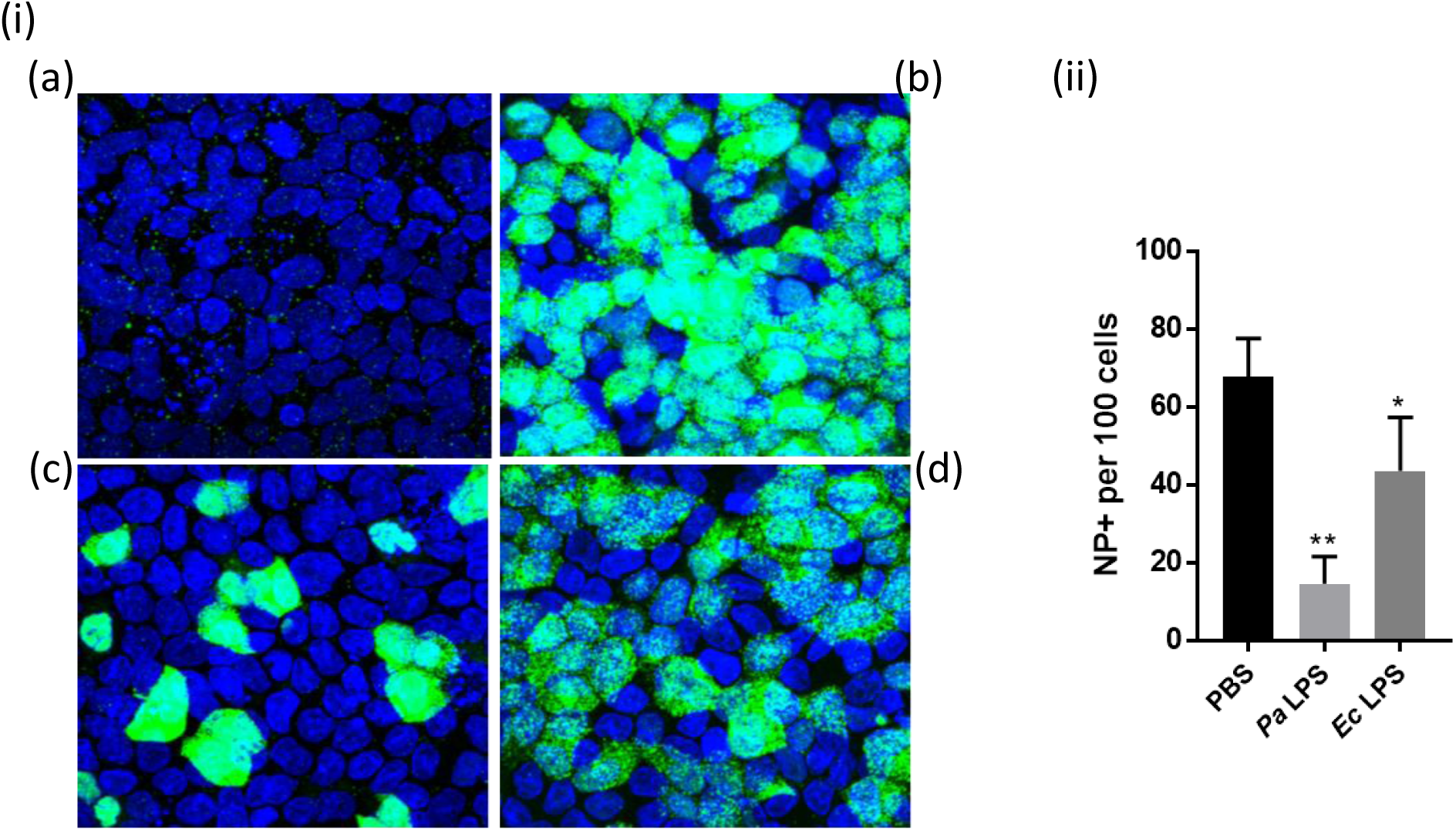
Reduction of infectivity by IAV following incubation with LPS. IAV diluted to an MOI of 2 were co-incubated at 33°C for 2 hours with PBS or LPS derived from *P. aeruginosa* or *E. coli*., then immediately used to infect MDCK cells. Cells were fixed at 7h post infection and stained with antibody against the viral nucleoprotein (green). Nuclei were stained with DAPI (blue). (i) Representative images of uninfected cells (a), cells infected with Neb/14 in PBS (b), cells infected with Neb/14 coincubated with LPS from *P. aeruginosa* (c), and cells infected with Neb/14 coincubated with LPS from *E. coli* (d) are shown here. n/a: no cells were infected. (ii) The data are representative of the mean NP+ cells and standard deviation obtained from two independent replicates, each with two technical replicates. Statistical significance relevant to Neb/14 incubated in PBS (control), was determined by one-way analysis of variance (ANOVA). *p<0.02, **p<0.0001

Next, we investigated total reductions in infectious virus following incubation of IAV with LPS derived from *P. aeruginosa* and *E. coli* at 33°C and 42°C, temperatures emulative of the human URT and the avian enteric tract (Figure 8). Since LPS is known to be an immunogenic stimulator, we first ascertained if incubating cells with LPS prior to infecting with IAV would confound the results of our assay. Cells incubated with LPS prior to infection yielded similar levels of infectious virus titers compared to cells incubated with LPS and IAV together (data not shown). In agreement with data obtained from IAV internalization assays using the viral NP protein shown in Figure 7, coincubation with *E. coli* resulted in no statistically significant loss of infectivity observed for any IAV tested relative to untreated controls at both temperatures. In contrast, incubation of IAV with LPS from *P. aeruginosa* resulted in greater reduction of virus infectivity compared to incubation with LPS from *E. coli*. Additionally, incubation of IAV with LPS from *P. aeruginosa* caused a greater loss of infectivity at 42°C than at 33°C for human and avian IAV. Our results suggest that LPS derived from *P. aeruginosa* and *E. coli* differentially impact retention of infectivity of human A(H1N1)pdm09 IAV.

**Figure 8.**
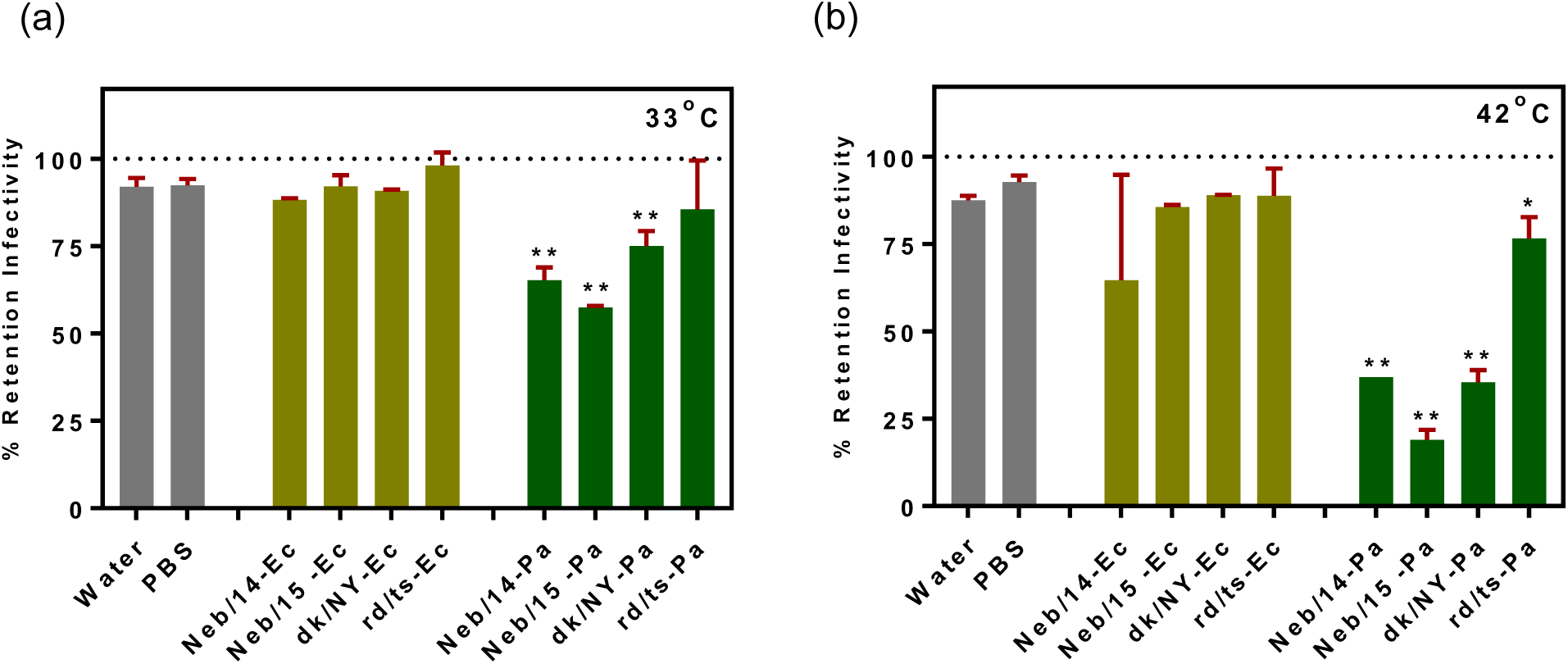
Bacterial LPS mediated effect on infectivity of human and avian IAV. Infectivity of human and avian IAV after incubation with LPS from *E. coli* (Ec) and *P. aeruginosa (Pa)* for 2 hours at (a) 33°C and (b) 42°C are shown here. 100% infectivity, indicated by the dotted line, was obtained by incubating IAV in PBS at 4°C and used as a control. Homologous virus was obtained by incubating all IAV in water or PBS at the two temperatures and taking the mean percent retention of infectivity across all IAV. The amount of viable virus in each treatment condition at the two temperatures, after each assay was determined by standard plaque assay, and compared to that in a 4°C PBS control to calculate the percent retention of infectivity. The data are representative of the mean and standard deviation of three independent replicates. Statistical significance relevant to 100 percent infectivity indicated by the dotted line, was determined by one-way analysis of variance (ANOVA). *p<0.02, **p<0.005

## Discussion

Person-to-person spread of IAV (during annual epidemics and occasional pandemics) is predominantly mediated by expulsion of virus-laden aerosols generated from viral replication occurring in the human URT. This host niche is also occupied by a wide diversity of commensal and pathogenic bacterial species: IAV can directly contact these bacteria and their surface structures before attaching to and infecting host cells. Previous research investigating the nature of these interkingdom microbial interactions between influenza viruses and host bacteria has largely used human seasonal viruses (10, 13, 29), which are already well-adapted to replicate in the presence of these bacterial species. Despite the wide diversity of host range determinants governing introduction of zoonotic IAV to humans, specific investigation has not been conducted into the potential barriers avian-origin IAV must overcome to successfully maintain an infectious phenotype in a host niche containing microbial species not previously encountered by the virus. Our study provides side-by-side comparisons of both human and avian origin A(H1N1) viruses to specifically examine if IAV that are not well adapted to the mammalian URT are able to maintain an infectious phenotype in a non-niche environment. Furthermore, while this study is limited to human and avian A(H1N1) viruses, it contributes to a knowledge base examining virus-host bacteria interactions with other A(H1N1) strains, such as variant viruses from swine populations (30), and different IAV subtypes present in zoonotic reservoirs.

Consistent with previously published work (31), we observed similar growth kinetics of our human strains, Neb/14 and Neb/15, in Calu-3 cells at both 33°C and 37°C (Figure 2) independent of subtle differences in amino acid residues in the viral HA (Table 1). However, replication kinetics of the avian strains, dk/NY and rd/Ts, exhibited divergent replication capabilities between each other at both temperatures (Figure 2) despite having the same avian consensus sequence in the HA (Table 1). These differences could be attributable to other internal genes such as the viral polymerases which have a known role in IAV replication (32). The capacity of dk/NY to reach comparable titers to the human adapted viruses in a human respiratory tract cell line demonstrates the need to examine the replicative fitness of zoonotic IAV in mammalian cells, including in epithelial cells derived from the nasal passages which predominantly express α-2,6 linked SA (33, 34). Our findings are suggestive of the potential that these non-human host origin viruses might pose a public health threat even when maintaining an α-2,3 SA binding preference.

IAV encounter a plethora of bacterial species in their different host niches. Prior work has shown that IAV are able to bind to the surface of both Gram negative and Gram positive bacterial pathogens of the human respiratory tract including *P. aeruginosa*, *Moraxella catarrhalis*, *S. pneumoniae* and *S. aureus* to different degrees; bacterial pathogens bound to IAV demonstrated increased adherence to respiratory epithelial cells (10). However, this prior study was performed using human IAV strains and did not address how interactions between IAV and bacteria of the host URT could modulate viral infectivity. Our study utilizes both human and avian IAV strains and a bacterial pathogen of the respiratory tract to assess (using a co-sedimentation assay approach) if viruses of different host origins could associate with the bacterium. We performed coincubation experiments for durations ≤2 hours, to emulate the short interval IAV may be present in the URT before binding to susceptible epithelial cells, finding that co-sedimentation of IAV with whole bacteria was possible within this window. Our results indicate that both human and avian IAV are capable of associating with *P. aeruginosa* in conditions emulative of the human URT, albeit to different degrees, with notable heterogeneity present not only across influenza viruses originating from different hosts, but also in viral strains from the same host species. Interestingly, coincubation of virus with this live bacterium did not result in a loss of infectivity for any IAV tested under the experimental conditions employed, though further research is warranted to better investigate the specific nature of the associations detected and the extent to which strain-specificity may contribute to these interactions. Our work is in agreement with prior studies showing that IAV can associate with bacteria present in the mammalian URT, while emphasizing that these viral-bacterial interactions may be highly specific to the IAV strain employed, warranting caution when contextualizing results across diverse viral strains. We chose *P. aeruginosa* for these assessments because it is a well-known pathogen that colonizes and infects the human respiratory tract (35). *P. aeruginosa* is also relatively understudied in inter-kingdom interactions between influenza viruses and bacteria of the host URT.

Since IAV encounter a multitude of bacterial species and their surface polysaccharides upon introduction into the human URT (27), we used microbial glycan microarrays to evaluate potential interactions between whole IAV derived from both human and avian hosts, and bacterial polysaccharides derived from both mammalian respiratory tract and extrapulmonary sites. These microarrays have been utilized previously to assess interactions between host immune factors and glycans derived from both Gram-negative and Gram-positive bacteria to evaluate specific recognition of microbial antigens followed by induction of host innate and adaptive immunity (28). To our knowledge, this is the first time that microbial microarrays have been used for the study of virus-bacteria interactions. We identified IAV strain-specific differences in binding to diverse polysaccharides that appeared to be independent of SA binding preference, though heightened commonality was observed when array targets were limited to polysaccharides derived from bacteria present in the mammalian respiratory tract. It is plausible that binding differences observed at the highest and lowest intensities were mediated, at least in part, by structural components present on these glycans. For example, the strongest signals were obtained from IAV binding to bacterial glycans on the array that had a terminal Glc-Nac or a heptose or rhamnose moiety. Collectively, similar patterns of binding observed between LPS from respiratory tract bacteria and diverse IAV supports a degree of conservation in IAV-bacteria interactions among human and zoonotic IAV, though there is a need to examine interactions beyond the polysaccharides included in the array.

Although the microarray allows us to investigate interactions between bacterial surface structures and IAV in a high-throughput manner, it cannot delineate what specific viral components mediate binding between these IAV and these glycans. While the viral HA is the main surface glycoprotein responsible for binding to host cell receptors, neuraminidase (NA), the other surface glycan for influenza A viruses, is also known to exert receptor binding properties, albeit to a lesser degree (36–38). Future studies could utilize purified viral recombinant HA instead of whole IAV to understand if the viral HA facilitates the binding to the bacterial glycans on the array. Another limitation of this format is that purified glycans printed on the array may not fully recapitulate the way these carbohydrates are naturally presented on microbes. To address this, we chose LPS isolates of *P. aeruginosa* and *E.coli* from the array, bacterial species that inhabit distinct host niches, for studies assessing functional interactions with IAV.

TEM analysis of interactions between IAV and LPS revealed direct binding interactions within 1 hour of coincubation. We also observed disruption of the surface envelope of Neb/14 virions at 33°C upon incubation with LPS derived from *P. aeruginosa* (Figure 6). Our TEM data are in agreement with our assessment of viral internalization following coincubation with LPS showing reduced infectivity within the first cycle of viral entry into cells (Figure 7). Taken together, our data support that IAV can interact with both bacteria and their surface components rapidly post-entry into environments assumed to emulate the nasal cavity. As supported by Figure 4, heterogeneous IAV can bind to a diversity of bacterial polysaccharides. However, bacterial polysaccharides exhibit structural diversity (39–42), and the functional consequences of these interactions may vary widely, including but not limited to considerations of viral stability examined here. Further study of interactions between IAV and other bacteria or their surface glycoproteins is needed to better understand whether our results of the association between IAV and *P. aeruginosa* are unique to this specific bacterium, as it suggests the potential for both synergistic and antagonistic interactions among IAV in well-suited niches with resident bacteria in these locations. Future work to understand interactions between influenza viruses and whole microbes on a broader scale could utilize whole microbe microarrays as used in previous research (43–45).

Previous research involving virus-bacteria interactions have shown that incubation of enteric viruses (including mammalian orthoreovirus, poliovirus, coxsackievirus and murine norovirus) with bacteria and their surface components imparted stability to virions, particularly at elevated temperatures, resulting in enhanced infectivity for these viruses (46–48). However, a study by Bandoro and Runstadler assessing influenza-bacteria interactions observed reduced stability of human and avian IAV when exposed to GI tract bacterial isolates and LPS from *E. coli* in a temperature-dependent manner (12). To evaluate if influenza viruses could derive a fitness benefit from synergistically interacting with bacterial components of the respiratory tract, we incubated IAV with LPS from *P. aeruginosa* and *E. coli*, common bacterial species of the respiratory tract and the GI tract, respectively. We observed lower infectious titers when IAV was incubated with LPS from *P. aeruginosa* compared to incubation with *E. coli*. Of interest, seasonal IAV that are well adapted to replication in the human respiratory tract, were impacted negatively in this assay by LPS from a respiratory tract bacterium, and this suggests different mechanistic interactions than those observed with enteric viruses. In contrast to the previous study (12), the IAV tested in our assay did not exhibit reduced infectivity following coincubation with *E. coli*, suggestive of the high potential for strain specific interactions with these polysaccharides. The diversity of IAV binding strength identified in our MGM to diverse LPS isolated from the same bacterial species, coupled with our strain-specific results identified in LPS-coincubation functional assays, underscores that these IAV-polysaccharide interactions can be highly strain-specific. As such, it is important to use caution when summarizing the consequences of IAV-bacterial interactions and extrapolating results to diverse IAV not specifically tested in laboratory studies.

There has been a long-standing need to study influenza-bacteria interactions from a public health standpoint (5). While acute influenza virus infection has long been recognized as a significant public health concern worldwide, subsequent co-infection with various bacteria can significantly increase the mortality rate of patients. Previous research targeted at understanding these interkingdom interactions has primarily been focused on the impact that influenza viruses have on host bacteria (with a particular focus on *S. pneumonia*), with reduced investigation of the functional consequences that bacteria in the respiratory tract might be exerting on these viruses. Most prior studies in this area have been restricted to human seasonal influenza strains and have not considered viral-bacterial interactions as a potential host range determinant of IAV. This study provides a bridge to the existing research by including both human and avian IAV, highlighting the heterogeneity that exists across influenza viruses originating from the same host species as well as the complexity that arises from reservoir spillover events. The increased spillover events of IAV from non-human reservoirs to human populations, as currently evidenced by ongoing sporadic detection of A(H5N1) virus infection in humans in North America, highlights the need to study these interkingdom interactions between IAV and host niche specific bacteria (49). Influenza-bacteria coinfections are significant in the context of One Health, which emphasizes the interconnectedness of human, animal, and environmental health (50). Surveillance of these interactions in both human and animal populations can help identify emerging threats, enabling timely public health responses. By recognizing the complex relationships within ecosystems, we can better address the challenges posed by infectious diseases, ultimately improving health outcomes across all domains.

## Acknowledgements

Authors thank Dr Richard Webby (St Jude Children’s Research Hospital) for access to the A/ruddy turnstone virus used in this study, Dr. Lauren Byrd-Leotis and Dr. David Steinhauer (Emory University) for their help with virus purification and labeling, and Dr. Xuezheng Song and Yi Lasanajak (Emory University Glycomics Core) for helpful conversations about the MGM arrays. The findings and conclusions in this report are those of the authors and do not necessarily represent the official position of the Centers for Disease Control and Prevention and the Agency for Toxic Substances and Disease Registry.

